# EXTRACELLULAR PERINEXAL SEPARATION IS A PRINCIPAL DETERMINANT OF CARDIAC CONDUCTION

**DOI:** 10.1101/2023.05.25.542366

**Authors:** William P. Adams, Tristan B. Raisch, Yajun Zhao, Rafael Davalos, Sarah Barrett, D. Ryan King, Chandra B. Bain, Katrina Colucci-Chang, Grace A. Blair, Alexandra Hanlon, Alicia Lozano, Rengasayee Veeraraghavan, Xiaoping Wan, Isabelle Deschenes, James W. Smyth, Gregory S. Hoeker, Robert G. Gourdie, Steven Poelzing

## Abstract

**Rationale:** Cardiac conduction is understood to occur through gap junctions. Recent evidence supports ephaptic coupling as another mechanism of electrical communication in heart. Conduction via gap junctions predicts a direct relationship between conduction velocity (CV) and bulk extracellular volume. Ephaptic theory is premised on the existence of a biphasic relationship between CV and the volume of specialized extracellular clefts within intercalated discs.

**Objective:** Determine the relationship between ventricular CV and structural changes to micro and nano-scale extracellular spaces.

**Methods:** Conduction and connexin43 (Cx43) gap junction protein expression were quantified from optically mapped guinea pig whole-heart preparations perfused with albumin, mannitol, dextran 70kDa, or dextran 2MDa. Peak sodium current was quantified from isolated guinea pig ventricular myocytes. Extracellular resistance (R_e_) was quantified by impedance spectroscopy. Intercellular communication was assessed in a heterologous expression system with fluorescence recovery after photobleaching. Perinexal width was quantified from transmission electron micrographs.

**Results:** CV was significantly reduced by mannitol, and increased by albumin, dextran 70kDa and 2MDa. The combination of albumin and dextran 70kDa decreased CV relative to albumin alone. R_e_ was reduced by mannitol, not significantly changed by albumin, and increased by both dextran 70kDa and dextran 2MDa. Cx43 gap junction expression and conductance, and peak sodium current were not significantly altered by the osmotic agents. The perinexal width in response to osmotic agents, in order of narrowest to widest, was: albumin with dextran 70kDa, albumin or dextran 2MDa alone, dextran 70kDa or no osmotic agent, and mannitol. When compared in the same order, CV was biphasically related to perinexal width.

**Conclusions:** Cardiac conduction does not correlate with bulk tissue impedance, but is biphasically related to perinexal separation, providing evidence that the relationship between CV and extracellular volume in ventricular myocardium is determined by ephaptic mechanisms under conditions of normal gap junctional coupling.

## INTRODUCTION

Understanding cardiac conduction is critical to the diagnosis, treatment, and prevention of arrhythmogenic diseases of the heart. Cardiac conduction is traditionally understood to follow a cable-like model of electrical conduction.^1–4^ In this paradigm, electrical current is envisaged as flowing into myocytes, through gap junction (GJ) channels at the intercalated discs (ID), back across cell membranes through outward potassium channels, and returning via the extracellular space. Importantly, experimental evidence has provided support for this long-established theoretical conception by demonstrating decreasing extracellular resistance can increase conduction velocity (CV).^2, 5, 6^

Since electrical propagation occurs in the hearts of lower vertebrate animals in which GJs are scarce,^7–9^ and persists in mammalian hearts with the principal GJ protein connexin43 (Cx43) knocked out,^10^ computational models of conduction were revised to propose an additional method of electrical intercellular communication.^11–16^ In these models of ephaptic coupling (EpC: non-GJ, non-synaptic coupling), strongly depolarizing inward currents facing a 10-30nm cleft separating adjacent cardiomoycytes significantly decrease the extracellular potential to depolarize nearby membranes. In the case of dense sodium channel localization to the ID between cardiomyocytes, and more specifically in the GJ adjacent perinexus,^17^ sodium channels self-*activate* in a positive feedback process that rapidly decreases the extracellular potential and activates more sodium channels^18^ while simultaneously increasing intracellular potentials. Importantly, the rapid decrease of extracellular potentials, increase in intracellular potentials, and depletion of extracellular sodium in the cleft,^13, 19^ decreases the driving force during intercellular propagation in a process known as self-*attenuation*.^12^ All computational models with EpC reveal that starting from a very narrow cleft, widening clefts between cardiomyocytes will first *increase* CV by reducing self-attenuation and then *decrease* CV by reducing self-activation. Importantly, models with EpC predict the relationship between CV and cleft width is biphasic when GJ conductance is closer to experimentally measured values of gap junctional coupling (GJC).^20–27^

In order for conduction to occur, current must return to depolarizing myocytes through the bulk extracellular space, and cardiac CV should be directly proportional to extracellular volume if conduction were only dependent on cable-like propagation through GJs without EpC. However, we previously demonstrated that cardiac CV can be inversely correlated to extracellular volume.^28^ Subsequent studies revealed that the osmotic agent mannitol, which increases bulk extracellular volume, also increases perinexal volume and reduces CV.^29^ Since we found that ventricular conduction slowed with perinexal expansion,^30, 31^ we concluded that EpC is a significant determinant of CV. However, the relative contributions of cable-like propagation through bulk interstitial volume and ephaptic-like conduction through perinexal domains remain unresolved. The purpose of this study was to determine the relationship between ventricular CV and structural changes to bulk extracellular volume and perinexal nanodomains. Results demonstrate osmotic agents alter CV primarily via an ephaptic mechanism.

## METHODS

All animal research for this study was carried out in accordance with the principles of the Basel Declaration and recommendations of the Guide for the Care and Use of Laboratory Animals, National Research Council. The protocol was approved by the Institutional Animal Care and Use Committee (IACUC) at Virginia Polytechnic Institute and State University (Protocol #21-091)

### Data Availability

Raw data are available upon request. Details of all statistical methods and experimental materials are available in supplement material (Supplemental Statistics and the Major Resource Table respectively).

### Guinea Pig Langendorff Preparations

Adult male Hartley guinea pigs from Hilltop Lab Animals, Inc. (13-15 months; 900-1200 grams) were chosen for these experiments in order to compare positive control to previous ephaptic studies using the same age, sex and species.^28, 29^ Animals were anesthetized by isoflurane inhalation. and hearts were excised and quickly (<5 minutes) cannulated in a Langendorff configuration using an oxygenated perfusion solution containing, as previously described,^28^ (in mmol/L): 1.25 CaCl_2_•2H_2_O, 140 NaCl, 4.5 KCl, 10 Dextrose, 1 MgCl_2_•6H_2_O, 10 HEPES, 5.5 mL/L of 1N NaOH and the electro mechanical uncoupler 2,3-butanedionemonoxime to reduce motion. The perfusate was equilibrated to a pH of 7.41 at 37°C using NaOH or HCl, as necessary. Hearts were immersed in a 3D printed poly(lactic acid) bath containing the perfusion solution maintained at 37°C.^32^ In all experiments, control solution was perfused for 30 minutes, followed by the same solution with either mannitol (Sigma-Aldrich M4125, D-Mannitol, 26.1 g/L), albumin (Sigma-Aldrich A9647, Bovine Serum Albumin, 4 g/L), dextran 70kDa (Sigma-Aldrich 31390, dextran from *Leuconostoc* pp. M_r_ ∼70,000, 40 g/L) or dextran 2MDa (Sigma-Aldrich D5376, dextran from *Leuconostoc mesenteroides* average mol wt 1,500,000-2,800,000, 40 g/L) for 15 minutes. In preparing the osmotic agent solutions, dextran 2MDa was shaken to dissolve into solution. Mannitol, albumin, and dextran 70kDa concentrations were chosen for comparisons to previous studies^6, 28^ and dextran 2MDa was selected for its preferential restriction to vasculature.^33^

### Statistical Analysis

The total number of animals or plates used (N) and replicate measures (n) were determined in order to detect 20% effect size based on means and averages obtained in previous studies with similar techniques to achieve an alpha of 0.05 and power of 0.8. Differences among experimental groups were determined through ordinary one-way ANOVA, nested-ANOVA to account for experimental replicates within hearts or plates in heterologous culture experiments, and Student’s t-test, or with non-parametric tests when appropriate (Kruskal-Wallis test or Wilcox test). Corrections for multiple comparisons were applied within, but not across experiment types,;tests were carried out in GraphPad/Prism v9.0 or later. As the distribution of perinexal width (W_P_) has an elongated right tail, a generalized linear mixed effects model specifying a gamma distribution and identity link was performed using the lme4 package in R version 4.2.2 to compare test groups to Time Control. Applied statistical tests are noted in figure legends and details are provided in the supplement (Supplementary Statistics). Reported values are compared to baseline and Time Control experiments. Data in the text are presented as mean difference (Diff) with 95% confidence interval (CI) using the following notation (Diff: X units, 95%CI: ±Y). Figures include individual data points and the 25 to 75 percentiles with whiskers representing the range, unless otherwise noted in the legends. Representative images were chosen from the complete data set with the following subjective criteria: 1) The image provided a clear visual example of the type of data being collected. 2) The difference shown between treatment conditions was closest to the average in the data as supported by statistical analysis. 3) The data represented the general quality of imaging collected.

### Optical Mapping

Conduction velocity (CV) was quantified by optical voltage mapping using di-4 ANEPPS (15 μM, Biotium #61010) as previously described.^28^ Briefly, the preparation was stained with di-4 ANEPPS by direct coronary perfusion for approximately 10 minutes after an initial 15-minute stabilization period following cannulation. Following a 20-minute dye wash-out period, the tissue was excited by 510 nm light, and the emitted light, filtered by a 610nm filter, was captured by a Micam Ultima L-type CMOS camera with 100×100 pixels and a magnification of 0.63X, covering an area of 15.9mm by 15.9mm. Hearts were stimulated at a basic cycle length of 300ms with a 5ms wide cathodal pulse at 1.5X the minimum current required to excite the tissue through a monopolar Ag-Ag/Cl electrode.

Optical data were analyzed to quantify transverse and longitudinal CV (CV_T_ and CV_L_, respectively), action potential duration (APD),^34–36^ and maximum dispersion of repolarization – the greatest time difference in repolarization between the left, right, base, and apex quadrants of the imaging field.^37, 38^ Supplemental Figure S1 demonstrates how CV_T_ and CV_L_ are measured from optical maps.

As albumin degrades optical signals,^30^ six consecutive paced beats were signal averaged prior to analysis for hearts perfused with albumin. APD was quantified as the time between activation and 90% repolarization. Half of the experiments were performed by a second blinded experimentalist and conditions were randomized for the purpose of rigor and reproducibility. In these experiments, test solutions were prepared by one experimentalist who knew the condition being tested and were provided to a second experimentalist with non-identifying markers (1 and2, A and B, or similar) while the second experimentalist selected the animal to be used, and carried out the isolation, perfusion, and mapping with no knowledge of the condition being tested. Experimentalists were unblinded only after CV measurements were made.

### Western Blots

Left ventricular (LV) tissue samples were snap frozen at specific time points in the protocol and western blotting was performed as previously described.^39^ Briefly, the samples were homogenized in RIPA lysis buffer (containing 50mM Tris pH 7.4, 150mM NaCl, 1mM EDTA, 1% Triton X-100, 1% sodium deoxycholate, 0.1% sodium dodecyl sulphate, 2mM NaF, 200μM Na3VO34) supplemented with Roche Protease Inhibitor Cocktail (4693159001, Sigma-Aldrich). Protein concentration was determined by a BioRad DC protein assay and concentrations were normalized prior to analysis. Electrophoresis was performed to separate proteins which were then transferred to a PVDF membrane, blocked with 5% bovine serum albumin for 1 hour at room temperature and incubated overnight with a primary antibody against the principal ventricular GJ protein Cx43 phosphorylated at Ser368 (pCx43, 1:1000, #3511S, Cell Signaling Technologies), at 4°C. The membranes were then washed and incubated with secondary antibody (1:5000, Goat Anti-Rabbit HRP, ab6721, abcam) at room temperature for 1 hour. After washing, bound antibody was detected using West Pico Plus chemiluminescent substrate (Thermo Scientific #34579) and imaged using the Licor Odyssey Fc system. Membranes were stripped with ReBlot Plus according to manufacturer’s instructions, blocked in Odyssey Blocking Buffer (Licor #927-50000) at room temperature for 1 hour and incubated with primary antibodies against total Cx43 (1:5000, C6219 rabbit, Sigma Aldrich) and GAPDH (1:5000, 101983-284 mouse, VWR). Membranes were then washed and incubated with secondary antibodies for 1 hour (both 1:10,000, goat anti-rabbit IRDye 800CW and goat anti-mouse IRDye 680RD) and washed again. Membranes were again imaged using the Licor Odyssey Fc system to determine protein expression, which includes an internal background subtraction algorithm that accounts for residual florescence and a specific secondary antibody binding. Total Cx43 was normalized to GAPDH and pCx43 was normalized to total Cx43. Primary antibodies for total Cx43 and pCx43 are widely used commercially available antibodies that have been validated previously.^40, 41^ The full immunoblots and ladders of representative data are presented in Supplemental Figure S2, as are all full uncropped blots used in this study.

### Fluorescence Recovery After Photobleaching (FRAP)

Rat glioma C6 cells (ATCC® CCL-107TM) transfected to stably overexpress Cx43^42^ were plated at ∼1.5e5 cells on a 35mm dish with 20mm glass bottom inset (Cellvis D35-20-1.5-N) and cultured for 48 hours in DMEM/F12 media mixture with 10% FBS, as previously described.^43, 44^ After 48 hours growth, cells were rinsed with Ca^2+^ and Mg^2+^ free DPBS and then incubated for 1 hour in a CO_2_-free incubator with Invitrogen Opti MEM with 5% FBS and all treatments but the GJ inhibitor carbenoxolone (CBX). CBX was added after dye loading to avoid interference with dye loading. After a 60-minute equilibration period, cells were rinsed and loaded with 2.5 µM calcein-AM, dissolved in serum-free Opti-MEM, and incubated for 15 minutes without any treatment. Cells were then rinsed and returned to Opti-MEM with 5% FBS and the appropriate treatment, and incubated for 5-10 minutes before imaging.

The cells were imaged on a Leica TCS SP8 confocal microscope using a 488nm laser at 0.5% power, 256×265 pixel resolution, 700Hz scan speed, and a 7.0 Airy unit pinhole. Pre-bleach images (n=5) were collected, then cells were bleached at 100% laser intensity for 50 cycles (377ms cycle duration). Post-bleaching, one image was collected every 0.5 seconds for 300 seconds. During the imaging procedure, cells were maintained near 37°C through use of a heated stage.

FRAP was measured for 15-25 cells per plate as replicates. The minimum fluorescence value was subtracted from each cell, so recovery is calculated from zero and normalized to a percent recovery, where the pre-bleach intensity is 1. Recovery was then fit to the following function: 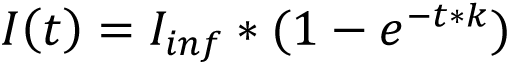.^44^ Coefficients *I_inf_* and *k* were calculated for each cell using the Matlab fit function with the nonlinear least squares method. Total number of plates per condition are as follows: control (N=13), carbenoxolone (N=6), mannitol (N=7), albumin (N=4), dextran 70kDa (N=4), and dextran 2MDa(N=4). All cells (replicates) were averaged, and the averages for each plate are treated and presented graphically as individual data points for each intervention. Relative recoveries (*I_inf_)* were statistically compared with GraphPad/Prism’s nested ANOVA function. The Nested parameter was the plate.

### Whole Cell Patch-Clamp Recording

Myocytes were isolated from LV free wall of guinea pig hearts using enzymatic dispersion technique as described previously^45^ and were re-suspended in 10 mL of Dulbecco modified Eagle medium, stored at room temperature, and used within 24 hours of isolation.

Sodium currents were recorded by ruptured-patch whole cell voltage clamp at room temperature. Microelectrodes were filled with a solution of (in mmol/L): CsF 120, MgCl_2_ 2, HEPES 10, EGTA 11 and brought to a pH of 7.3. Isolated myocytes were placed in the solution containing NaCl 25, N-methyl D-glucamine 120, CsCl 5, MgCl_2_ 1, NiCl_2_ 1, glucose 10, HEPES 10, pH 7.3. Sodium currents were elicited from a holding potential of -80 mV with depolarizing voltage pulses from -60 to 45 mV for 16 ms. Ionic current density (pA/pF) was calculated from the ratio of current amplitude to cell capacitance. Command and data acquisition were operated with an Axopatch 200B patch clamp amplifier controlled by a personal computer using a Digidata 1200 acquisition board driven by pCLAMP 7.0 software (Axon Instruments, Foster City, CA).

### Histology

A positive-pixel analysis of hematoxylin and eosin (H&E)-stained LV tissue, which had been fixed in 10% formalin for at least 24 hours, was performed to quantify interstitial volume (VIS), similar to what we have described previously.^28^ All measurements were made from the subepicardium, defined as the region from the epicardial surface to a depth of no more than 500 μm. Images of 5 μm-thick slices of stained tissue were analyzed by an Aperio positive-pixel analysis program which quantified interstitial space as a percentage of the entire array which was devoid of stained pixels.

### Extracellular Resistance

In a separate group of experiments, hearts were cannulated with the same perfusion solutions described above, but were not perfused with di-4-ANNEPS. Instead, after a similar stabilization period, a four-probe stainless steel electrode array, with the electrodes 2 mm apart and extending to a depth of 2 mm, was placed on the anterior surface of the LV, approximately parallel to the LAD, and connected to a Gamry Interface 1000 Potentiostat. Using the Gamry Framework electrochemical impedance program, tissue impedance was measured with the following parameters: Frequency sweep: 1-1E10 Hz; points/decade: 10; ACV: 0.2mV; DVC: 0; Area: 1cm^2^; Estimated Z: 100 Ω.

The data were then fit to an electrical circuit model of a resistor in parallel with a constant phase element (CPE) and intracellular resistance (Supplemental Figure S3). Values for the impedance were measured over the frequency band of 200 Hz∼1E6 Hz to estimate R_e_ and CPE.^46, 47^ Impedance of CPE is expressed by:

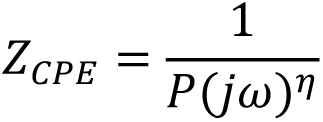

P and η are the parameters of CPE. At the tissue level, the cell membrane time constant is spatially distributed because of the variety of cells within the tissue. The distributivity of the time constant is described by the parameter η in CPE, which we can use as an index of the response of cell membrane electrical parameters to the addition of osmotic agents. After a baseline measurement, an osmotic agent was perfused for 15 minutes, followed by a final measurement. This 15 minute time point was selected to ensure we captured the impedance effects of osmotic agent perfusion observed in a previous study.^6^

### Fixable Dye Perfusion

In order to assess how different sized osmotic agents partition between the bulk interstitial volume and the ID, hearts were perfused with fixable dyes of different molecular weights: Alexa 488 hydrazide (0.5 kDa; ThermoFisher Scientific #A10436) and biotin-dextran (3 or 10 kDa; ThermoFisher Scientific, #D7135 and #D1956). After 30 minutes of dye perfusion, tissue samples collected from the anterior LV free wall were fixed in paraformaldehyde (2%) for 3 hours at 4°C, equilibrated sequentially in 15 and 20% solutions of sucrose, and frozen for cryosectioning. Sections were cut (5 µm) and immunolabeled as described above using a rabbit polyclonal anti-Cx43 antibody (Sigma-Aldrich C6219, St. Louis, MO) followed by goat anti-rabbit Alexa 568 (1:4000) and Streptavidin-conjugated Alexa 647 (1:4000) (ThermoFisher Scientific, #S21374) for biotin binding. Confocal imaging was performed using a TCS SP8 confocal microscope as previously described.^31^ Regions of interest were selected from confocal images surrounding IDs, which were identified based on Cx43 immunosignal. Fluorescence intensity of the green (0.5 kDa Alexa 488), red (3 or 10 kDa biotin-dextran), and magenta (Cx43) signals were averaged along an axis perpendicular to the ID to generate intensity profiles. The intensities of green and red signals at the maxima of the magenta (Cx43) signal were normalized to their respective median values on either side of the ID to yield an ID accumulation / depletion index. Values greater than 1 indicate accumulation within the ID while values close to 0 indicate exclusion from the ID.

### Electron Microscopy

Anterior epicardial tissue from the base of the LV (N=8 hearts x 5 interventions) was collected from hearts after the conclusion of the optical mapping experiments. Tissue was cut into 1 mm^3^ chunks which were fixed in 2.5% glutaraldehyde for at least 24 hours at 4°C before being washed and stored in phosphate-buffered saline (PBS) at 4°C. Samples were then prepared for transmission electron microscopy (TEM) and imaged as previously described using a JEM JEOL 1400 Electron Microscope at 150,000X magnification.^48^ To quantify W_P_, 15 images were analyzed per sample using a previously-described custom Matlab program.^48^ The average intermembrane distance between 30-105nm from the edge of the GJ plaque is reported as W_P_. Previous work measuring this parameter indicates that 10 to 15 images taken from a minimum of 4 hearts is sufficient to detect changes in W_P_ more than 2nm.^49^

## RESULTS

### Whole-Heart Electrophysiology

We quantified ventricular conduction velocity (CV) by optical mapping during perfusion with our historical lab standard solution^31^ (baseline) and after 15 minutes perfusion with the addition of one of the osmotic agents. In Figure 1A, representative optical maps demonstrate how CV is affected by time and osmotic agents. After 15 minutes, CV may modestly increase in time-control optical maps, is decreased with mannitol (as evidenced by crowding of isochrones), is increased by albumin (as evidenced by enhanced isochrone spacing), is modestly increased by dextran 70kDa, and increased by dextran 2MDa. Summary data in Figure 1B demonstrate the effect of the osmotic agents relative to baseline measures in the same heart as well as a comparison of the paired change in conduction to Time Controls. In short, CV in the transverse direction (CV_T_) was not significantly different after 15 minutes of Time Control perfusion (Diff: 1.1 cm/sec, 95%CI: ±1.3). Mannitol significantly decreased CV_T_ relative to baseline measurements (Diff: -2.7 cm/s, 95%CI: ±1.8), and the change in CV_T_ was significantly lower (Diff: -3.7 cm/s, 95%CI: ±3.0) than the change in CV_T_ observed in the 15-minute Time Control measurement. Albumin, on the other hand significantly increased CV_T_ (Diff: 7.2 cm/s, 95%CI: ± 2.4) relative to baseline, and the change was significantly greater (Diff: 6.1 cm/s, 95%CI:± 2.8) than the change found for the Time Control. Interestingly, both dextran 70kDa and dextran 2MDaincreased CV_T_ in paired comparisons (Diff: 1.0 cm/sec, 95%CI: ±0.9 and Diff: 2.7 cm/sec, 95%CI: ±1.6), but the change in CV_T_ was not significantly different (Diff: -0.1 cm/s, 95%CI: ±3.0 and Diff: 1.7cm/sec, 95%CI: ±3.0) from the Time Control. CV in the longitudinal direction (CV_L_) significantly increased after 15 minutes of Time Control perfusion (Diff: 4.1 cm/sec, 95%CI: ±3.9). Mannitol did not significantly change CV_L_ (Diff: -2.1 cm/sec, 95%CI: ±3.5). Albumin (Diff: 8.7 cm/sec, 95%CI: ±4.1) Dextran 70kD (Diff: 4.5 cm/sec, 95%CI: ±4.4) significantly increased CV_L_. Dextran 2MDa did not significantly change CV_L_ (Diff: 1.9 cm/sec, 95%CI: ±2.9). However, none of the magnitudes of change were significantly different from Time Control. In summary, epicardial transverse conduction relative to the paired pre-treatment baseline was decreased by mannitol, and increased by albumin and both dextrans.

**Figure 1.**
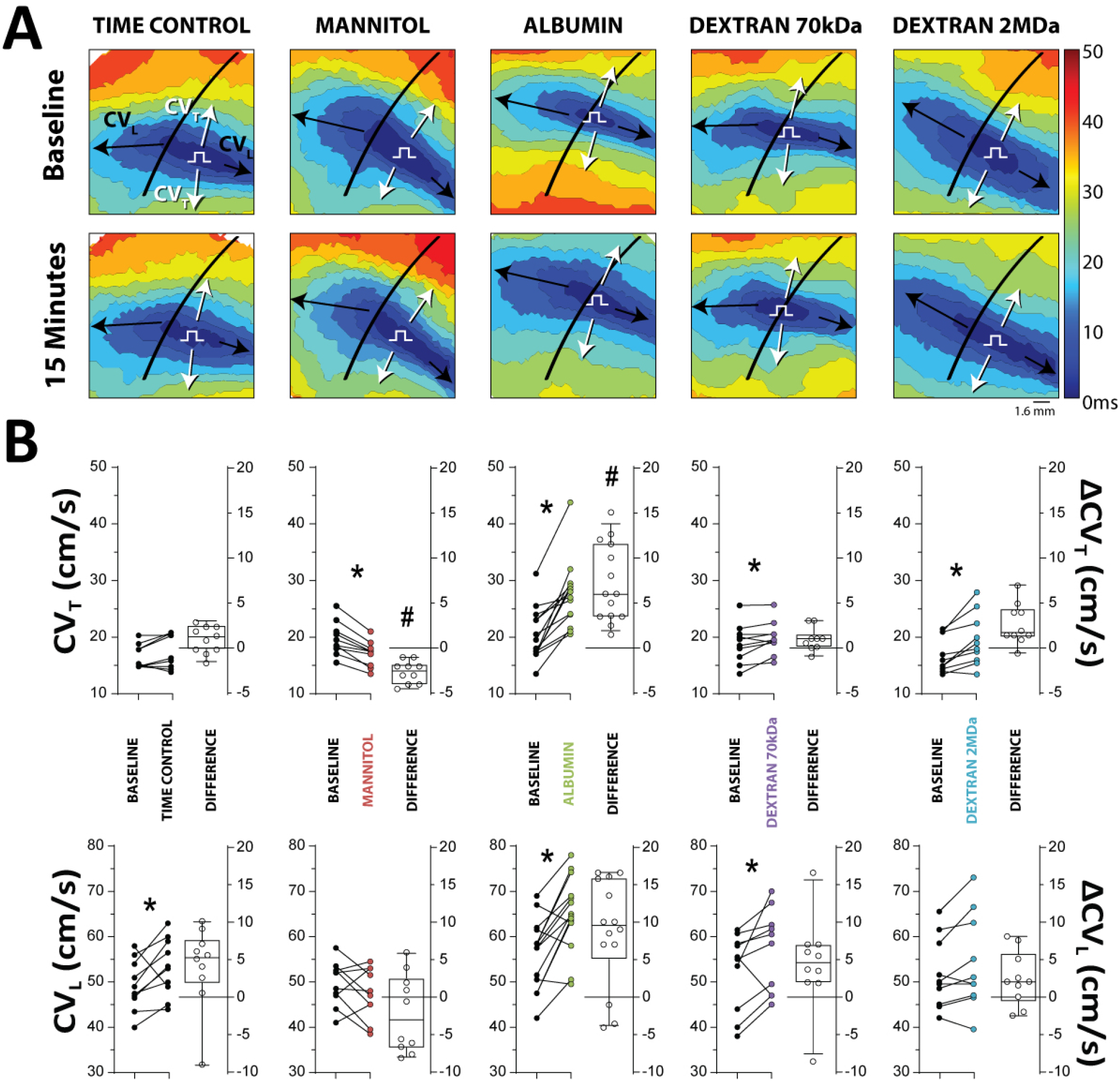
Osmotic agents modulate transverse conduction velocity. **A.** Representative 3ms optical mapping activation time isochrones demonstrate the effects various osmotic agents have on the elliptical spread of activation. Arrows indicate conduction velocity directions of interest for longitudinal (CV_L_-Black) and transverse (CV_T_-White) propagation, and П notes the approximate site of pacing in the field of view. The approximate location of the left anterior descending coronary artery is noted by a black line. **B.** CV plotted as individual points (*left* axis) obtained sequentially in the same heart and the change in CV (*right* axis) from paired baseline experiments demonstrates mannitol significantly decreases, albumin significantly increases, dextran 70kDa increases, and dextran 2MDa increases CV preferentially in the transverse direction (Time Control N=10, Mannitol N=10, Albumin N=14, Dextran 70kDa N=10, Dextran 2MDa N=10). *p<0.05 relative to baseline measurements made 15 minutes before intervention in the same heart, paired Student’s t-test. #p<0.05 relative to Time Control, ΔCV across groups compared with ordinary one-way ANOVA applying Dunnett’s correction.

To explore whether conduction changes might be attributable to more significant electrophysiologic remodeling, action potential duration (APD) and the maximum dispersion of APD among the left ventricular (LV) basal and right ventricular apical quadrants of the imaging field were compared as previously described.^50^ The osmotic agents did not alter APD or APD dispersion (Figure 2), suggesting that the mechanisms governing conduction changes observed are related to cell-to-cell electrical propagation, but not repolarization. Specifically, time-controlled APD measured at 0 and 15 minutes was not significantly different (Diff: 4.4 ms 95%CI: ±8.6). Mannitol did not significantly change APD (Diff: 3.4 ms, 95%CI: ±11.2), nor did albumin (Diff: 6.9 ms, 95%CI: ±11.0), dextran 70kDa (Diff: 3.5 ms, 95%CI: ±10.9), or dextran 2MDa (Diff: -1.4 ms, 95%CI: ±23.9). Relative to baseline, maximum APD dispersion was also not significantly different for Time Control (Diff: 3.6 ms, 95%CI: ±22.5), mannitol (Diff: 3.3 ms, ±22.5), albumin (Diff: 5.5 ms, 95%CI: ±19.8), dextran 70kDa (Diff: -1.8 ms, 95%CI: ±19.0), nor dextran 2MDa (Diff: -10.75 ms, ±22.1).

**Figure 2.**
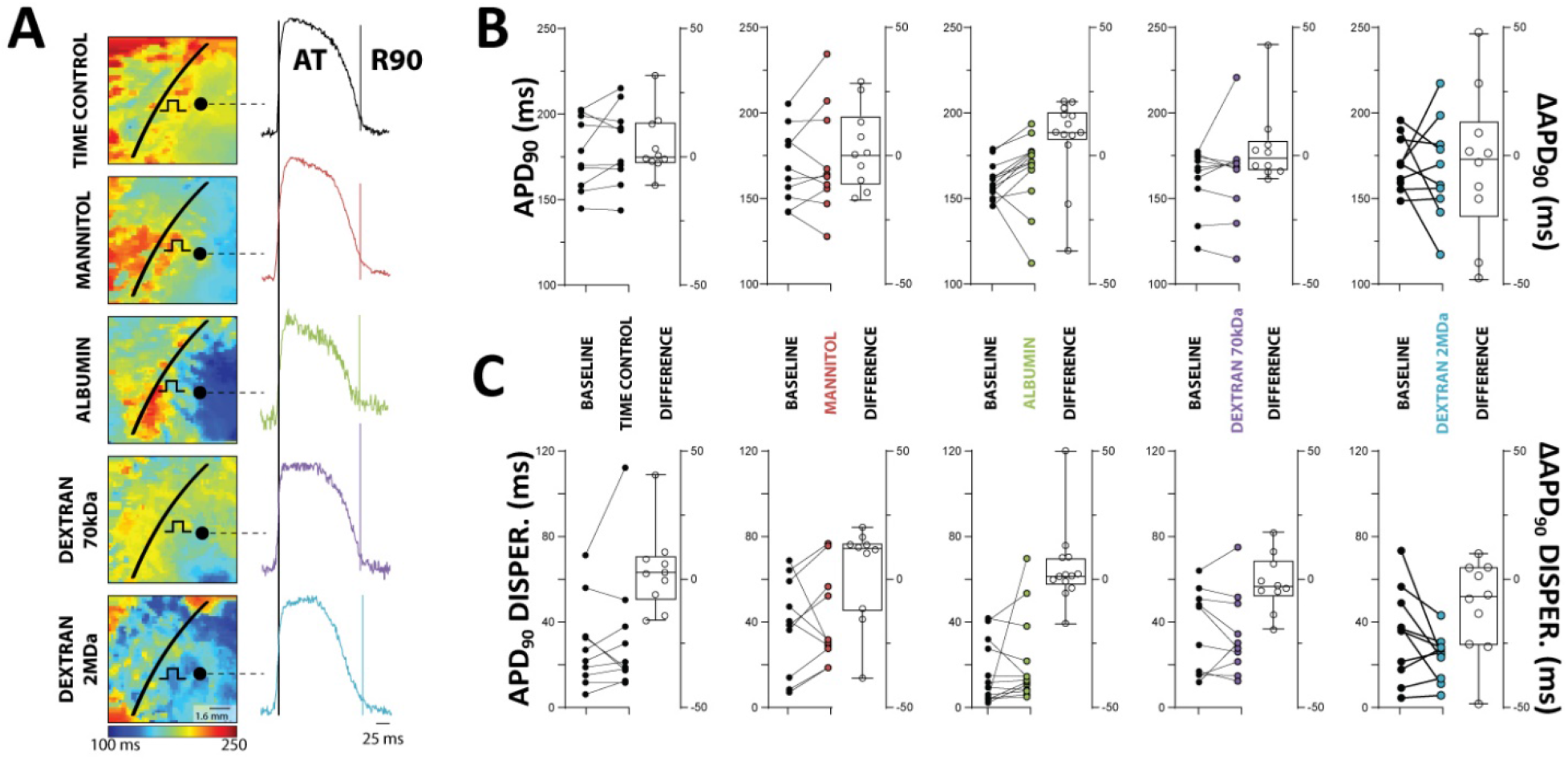
Osmotic agents do not change epicardial action potential duration (APD) in whole heart. **A.** Representative optical action potential duration (APD) maps and action potentials recorded from the anterior left ventricular epicardium. П notes the approximate site of pacing in the field of view. APD_90_ is the difference of Repolarization at 90% (R90) and Activation Time (AT). **B.** Summary APD_90_ data reveal that APD_90_ is not significantly affected by the osmotic agents utilized in this study, relative to pretreatment paired control, (paired Student’s t-test) nor are the magnitudes of the change different from that of the Time Control. p=n.s. ANOVA with multiple comparisons and Dunnett’s multiple comparisons test and Kruskal-Wallis test with Dunn’s multiple comparisons test) **C.** Furthermore, the maximum APD dispersion between quadrants of the imaging field are not significantly different from baseline (Time Control N=10, Mannitol N=10, Albumin N=13, Dextran 70kDa N=10, Dextran 2MDa N=10, nor were the magnitudes of change significantly different relative to Time Control. p=n.s, ordinary one-way ANOVA with Šídák’s multiple comparisons test and Kruskal-Wallis test with Dunn’s multiple comparisons test.

### Cx43 and Gap Junctions

In order to determine possible effects of osmotic agents on gap junctional coupling (GJC), levels of the principal ventricular gap junction (GJ) protein Cx43 were quantified by western blotting. Total Cx43 expression was not expected to change in response to 15 minute exposure to the osmotic agents, and so Cx43 phosphorylated at serine 368 (pCx43-S368) was also quantified since it has been associated with changes in GJ conductivity.^51–53^ Across all conditions, neither total nor pCx43-S368 (Figure 3A,B) significantly changed in response to osmotic agent perfusion. Relative to time-controlled tissue, total Cx43 was not significantly different in tissue perfused with mannitol (Diff: 0.1 AU, 95%CI: ±0.4), albumin (Diff: 0.3 AU, 95%CI: ±0.4), dextran 70kDa (Diff: -0.2 AU, 95%CI: ±0.4), nor dextran 2MDa (Diff: -0.02 AU, 95%CI: ±0.5). With respect to Time Control, pCx43-S368 was also not significantly different in tissue perfused with mannitol (Diff: 0.1 AU, 95%CI: ±0.6), albumin (Diff: -0.2 AU, 95%CI: ±0.6), dextran 70kDa (Diff: -0.1, 95%CI: ±0.6), or dextran 2MDa (Diff: -0.1 AU, 95%CI: ±0.5).

**Figure 3.**
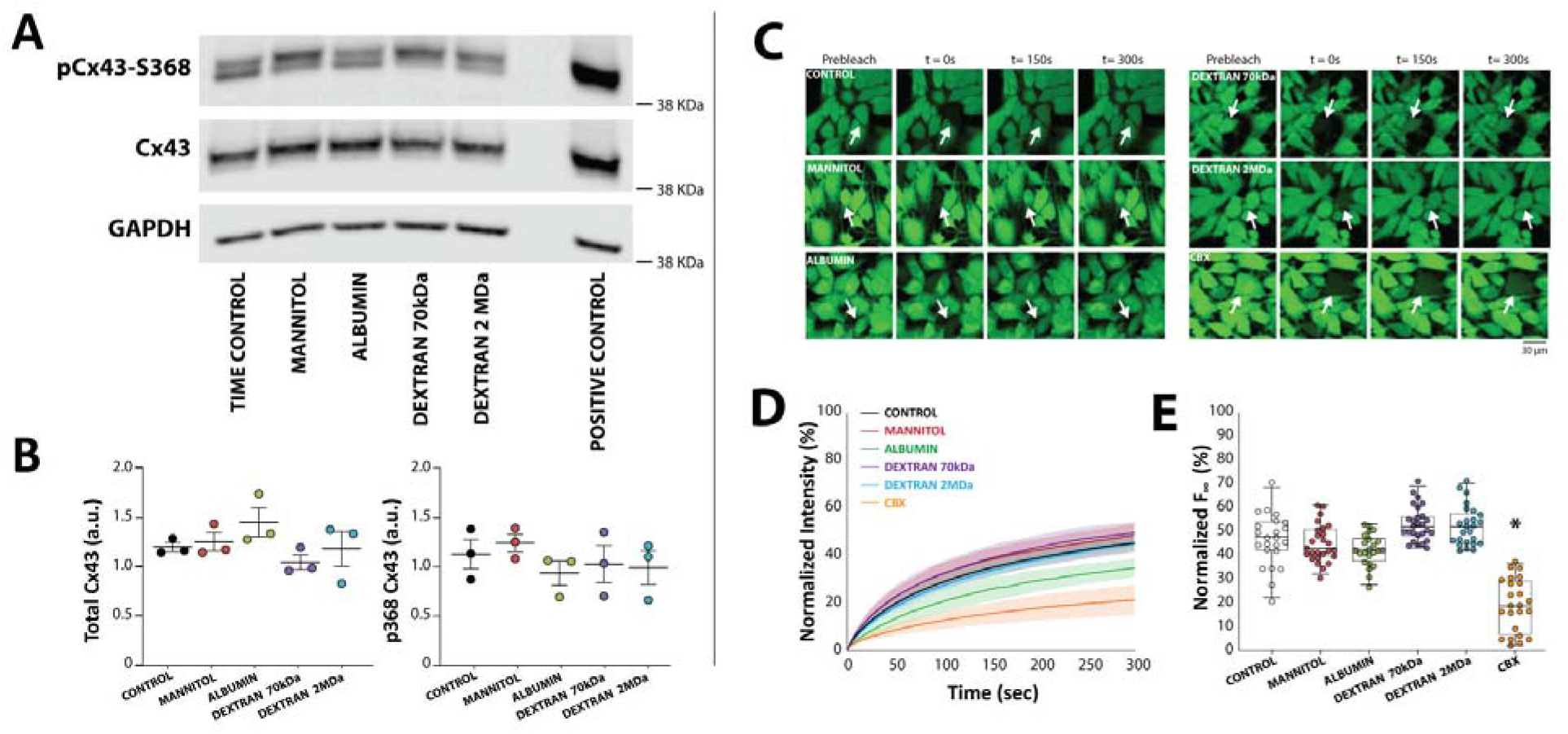
Osmotic agents do not significantly alter Cx43 protein expression or function. **A.** Representative western blots suggest that neither total connexin43 (Cx43) nor Cx43 phosphorylated at serine-368 (pCx43-S368) significantly change in response to acute exposure to osmotic agents. Lane 7 contains a positive control using hearts exposed to 60 min of room temperature ischemia, a condition shown to upregulate Cx43 pS368.^91^ **B.** Summary data plotted with mean and standard deviation error bars do not reveal significant differences in either total or pCx43-S368 for all conditions (N=3 hearts per condition. p=n.s. via ordinary one-way ANOVA applying Dunnett’s correction). **C.** Representative fluorescent images of calcein-AM fluorescence in control cultures (N=13 plates, n=220 cells) and cultures exposed to mannitol (N=7 plates, n=116 cells), albumin (N=4 plates, n=77 cells), Dextran 70kDa (N=4 plates, n=102 cells), Dextran 2MDa (N=4 plates, n=95 cells), and carbenoxolone (CBX, N=6 plates, n=110 cells). Measurements were made before bleaching (Pre-bleach), just after bleaching (t=0s), halfway through and at the end of the recovery period (t=150 and 300s, respectively). **D.** Fluorescence recovery after photobleaching (FRAP) plot of bleached cells during 300s post bleaching. Solid lines represent averages at each point, and shaded areas represent the standard error of the mean. **E.** Normalized recovery demonstrates that only CBX significantly decreased FRAP relative to control. *p<0.05, nested ANOVA applying Dunnett’s correction with culture plate nested under treatment type.

Since protein expression does not always correlate with function, functional GJC was also assessed by Fluorescence Recovery After Photobleaching (FRAP) in heterologous cells stably expressing Cx43 using an approach that we have reported previously.^43, 44, 54^ Representative fluorescent images taken before photo-bleaching (prebleach, Figure 3C), immediately after bleaching (t=0s), and at 150 and 300s post-bleaching, demonstrate fluorescence recovers to approximately the same extent as control cells. Specifically, FRAP, as can be seen in summary data plotted in Figures 3D and E, was not significantly different for cells incubated with mannitol (Diff: 1.0%, 95%CI: ±16.2), albumin (Diff: -4.2%, 95%CI: ±20.9), dextran 70kDa (Diff: 3.8%, 95%CI: ±20.7), or dextran 2MDa (Diff: 3.6%, 95%CI: ±20.8). As a positive control, the GJ uncoupler carbenoxolone (CBX, 10μM) decreased FRAP as expected (Diff: -23.8%, 95%CI: ±5.9). Despite a visual drop in FRAP measured in cells treated with albumin, summary data in Figure 3E reveal that only CBX significantly decreased GJC estimated by FRAP. Even if 1-hour perfusion with albumin modestly decreased GJC, that finding would be inconsistent with increased CV observed in hearts perfused with albumin. In summary, we did not measure a change in Cx43 level or function in response to the osmotic agents employed in this study at the concentrations or incubation times noted.

### Peak Sodium Current

Next, we sought to determine whether the osmotic agents affected the sodium current, which is another important determinant of cardiac conduction. Representative current traces from a single cell, the average of average current-voltage (IV) measurements from multiple cells in each heart, and summary peak current for all cells and hearts are plotted in Figure 4A-D. In summary, the osmotic agents did not significantly change peak sodium currents relative to control replicates obtained from the same hearts. Specifically, the peak sodium current relative to control was not significantly different in cells treated with mannitol (Diff: 4.1 pA/pF, 95%CI: ±11.0), albumin (Diff: -1.1 pA/pF, 95%CI: ±13.9), dextran 70kDa (Diff: -2.6 pA/pF, 95%CI: ±14.1), or dextran 2MDa (Diff: - 0.7 pA/pF, 95%CI: ±14.0).

**Figure 4.**
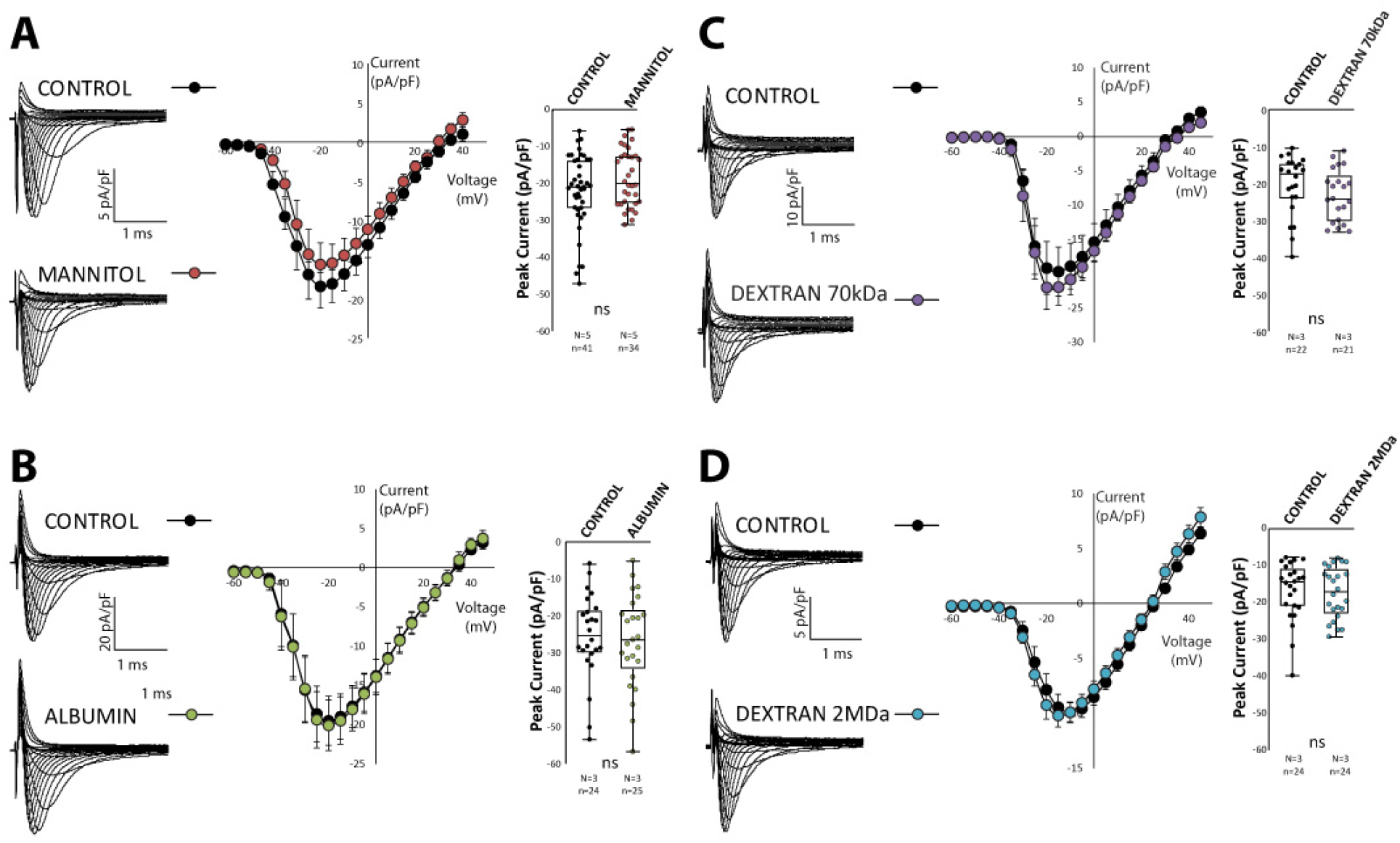
Osmotic agents do not significantly alter peak sodium current. **A.** Representative currents from a single cell, IV curves averaged from a single heart, and peak current at -20mV step potential for all hearts and cells reveal mannitol (N=5 hearts, n=34 cells, red) does not significantly decrease sodium currents relative to control measurements made in cells from the same heart (N=5 hearts, n=41 cells, black). **B.** Albumin (N=3 hearts, n=25 cells, green) did not significantly affect sodium currents relative to control (N=3 hearts, n=24 cells, black). **C.** Dextran 70kDa (N=3 heart, n=21 cells, purple) did not significantly affect sodium currents relative to control (N=3 heart, n=22 cells, black). **D.** Dextran 2MDa (N=3 hearts, n=24 cells, blue) did not significantly affect sodium currents relative to control (N=3 hearts, n=24 cells, black). All treatment groups were compared to same day measurements from control group cells using a nested ANOVA applying Šidák’s correction, where the heart is the nested variable.

### Bulk Interstitium

To test whether the volume of bulk interstitial space (VIS) correlated with changes in CV, the percentage of tissue unoccupied by cardiac cells was quantified with a positive pixel analysis of hematoxylin and eosin (H&E) stained tissue. Histological quantification did not reveal significant differences between tissues subjected to 15 minutes of perfusion with any osmotic agent (Figure 5). Relative to control, VIS was not significantly changed by perfusion with mannitol (Diff: 0.7%, 95%CI: ±14.1), albumin (Diff: 2.8%, 95%CI: ±14.6), dextran 70kDa (Diff: -6.1%, 95%CI: ±11.8), or dextran 2MDa (Diff: -1.2%, 95%CI: ±13.2). This was unexpected since our previous results from ventricular tissue analyzed from glutaraldehyde fixation and snap frozen samples demonstrated albumin decreases bulk extracellular volume and mannitol increases it.^28^ We therefore sought to quantify tissue impedance at baseline and after 15 minutes of treatment with the osmotic agents with a 4-electrode technique as previously described.^46, 47, 55^ Extracellular tissue resistance (R_e_) was measured in each heart without interventions (Time Control) and after serial perfusion with the osmotic agents. This permitted a pair-wise comparison to reduce errors associated with different depths of electrode penetration and orientation of electrodes relative to myocardial ultrastructure and transmural rotational anisotropy. The specific values of R_e_ are less important than whether the osmotic agent changed R_e_, and if R_e_ increased or decreased. Summary data in Figure 6A reveal that R_e_ did not significantly change over 15 minutes relative to baseline for Time Control measurements (Diff: 13.6Ω, 95%CI: ±89.7), mannitol significantly reduced R_e_ (Diff: -4.9Ω, 95%CI: ±5.0), albumin did not change R_e_ (Diff: 14.3Ω, 95%CI: ±85.7), dextran 70kDa significantly increased R_e_ (Diff: 20.6Ω, 95%CI:±10.9), and dextran 2MDa significantly increased R_e_ (Diff: 70.3Ω, 95%CI: ±68.4). Importantly, CV changes observed above do not correlate with R_e_.

**Figure 5.**
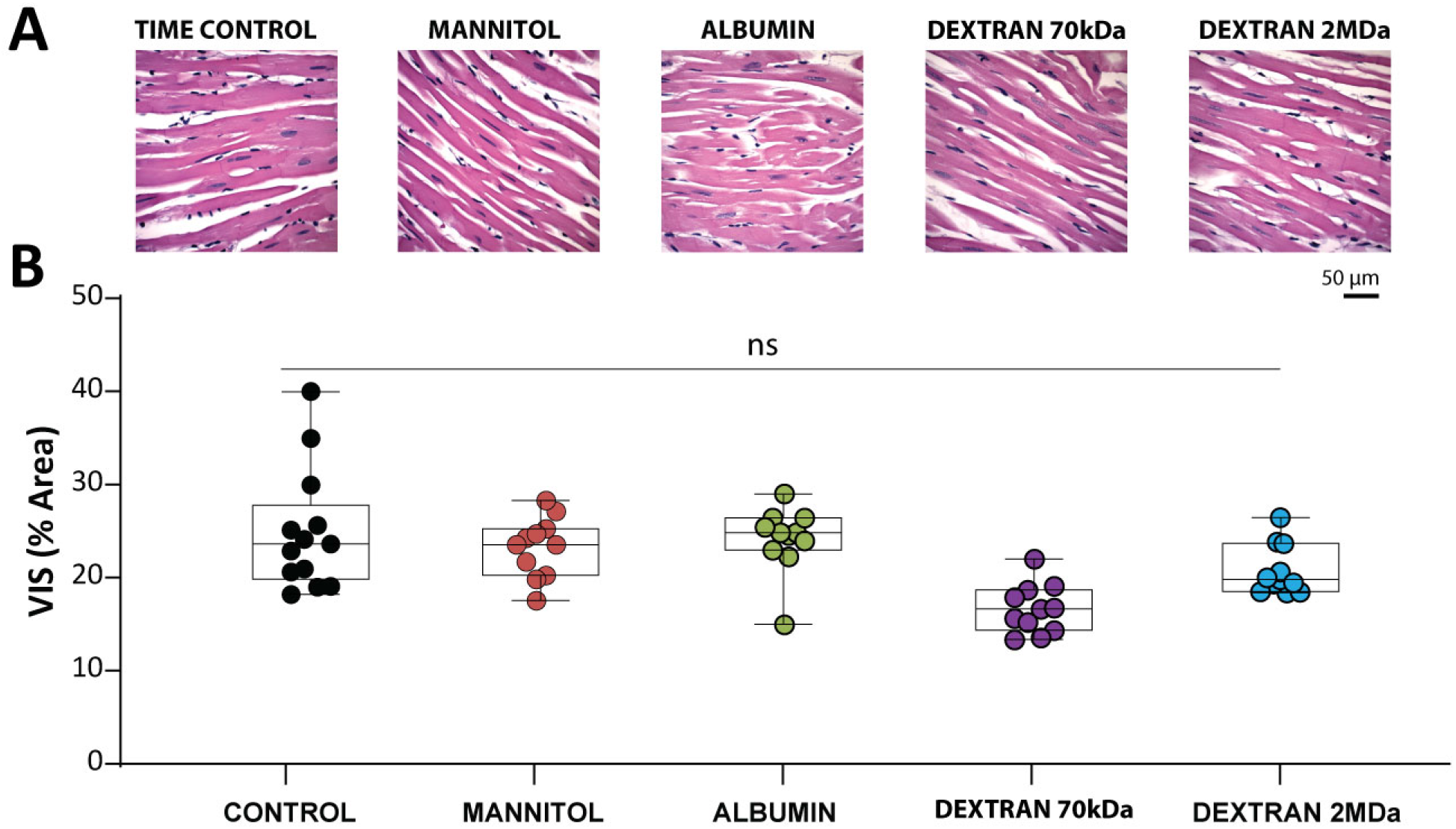
Analysis of H&E-stained ventricular tissue reveals no significant differences in interstitial space. The structure of bulk interstitium does not explain observed conduction changes via a cable-like mechanism, as demonstrated by **A.** representative images of H&E stained tissue with **B.** VIS quantified by a positive-pixel algorithm for hearts perfused with control (N=8 hearts, n=84 images), mannitol (N=7 hearts, n=73 images), albumin (N=7 hearts, n=71 images), dextran 70kDa (N=9 hearts, n=1 images), or dextran 2MDa (N=6, n=62 images). p=n.s. nested one-way ANOVA applying Šidák’s correction, with heart nested under treatment type.

**Figure 6.**
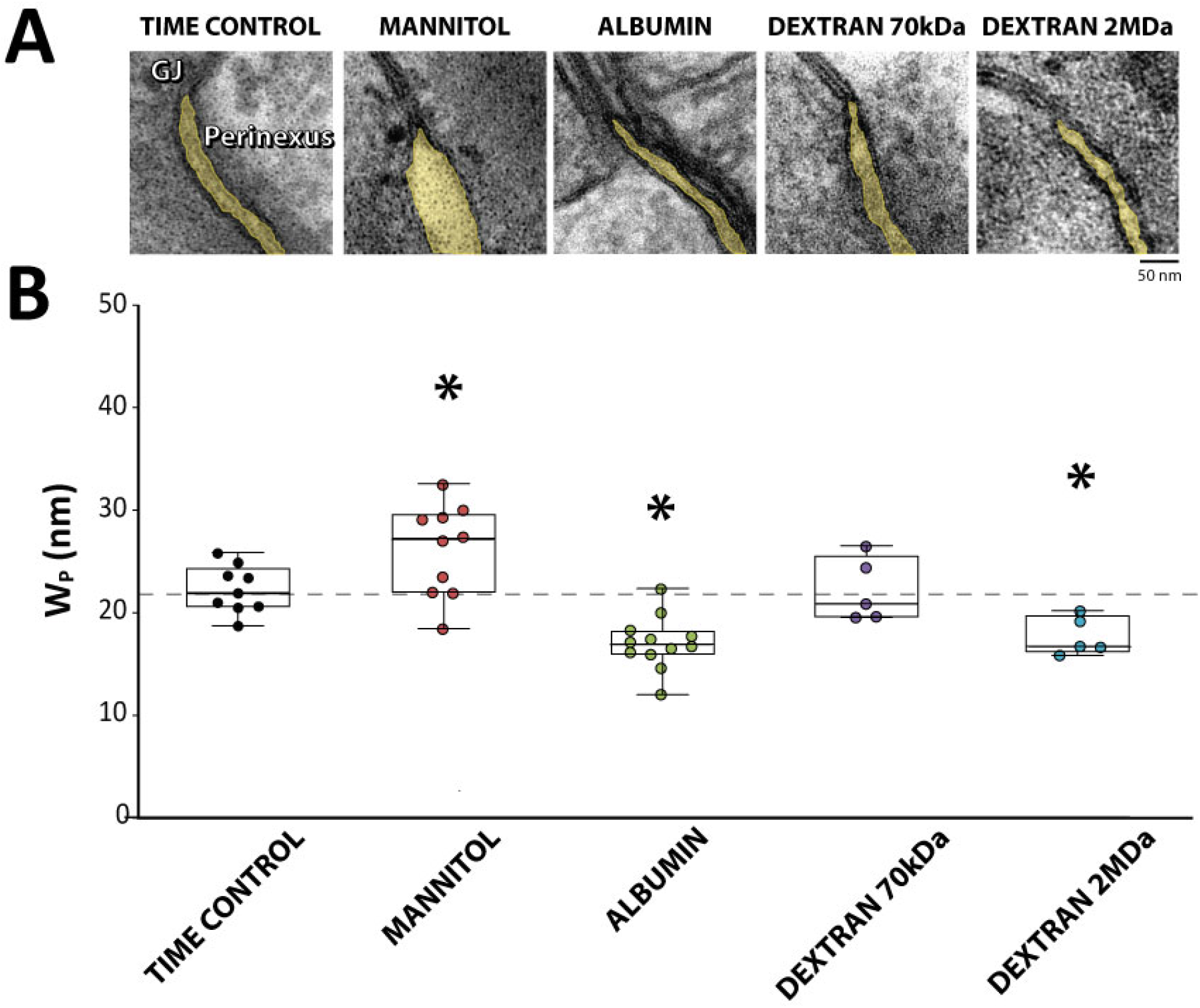
Osmotic agents modulate extracellular resistance. **A.** Neither extracellular resistance (R_e_), nor the constant-phase element (CPE) parameter η – an index of cell membrane electrical properties within intact tissue – varied during Time Control (N=4). **B.** Mannitol decreased R_e_ but did not change η (N=7). **C.** Albumin did not affect either R_e_ or η (N=4). **D.** However, Dextran 70kDa significantly increased R_e_ (N=4), but not η. **E.** Similarly, Dextran 2MDa increased R_e_ without altering η (N=4). *p<0.05, paired 2-tailed Student’s t-test, relative to Baseline.

Intercellular parameters, including cell membrane capacitance, cytosolic resistance and GJ resistance were modeled as a constant phase element (CPE). The distributivity of the time constant of the CPE element, the parameter η, did not significantly change relative to paired baseline measurements: Time Control (Diff: 0.05 AU, 95%CI: ±0.9), mannitol (Diff: 0.11AU, 95%CI: ±0.10), albumin (Diff: 0.00 AU, 95%CI: ±0.23), dextran 70kDa (Diff: -0.03AU, 95%CI: ±0.16), and dextran 2MDa (Diff: -0.03AU, 95%CI: ±0.20), as can be seen in Figure 6. These data suggest the R_e_ changes described above are unlikely to be the result of changes to cellular or tissue factors beyond R_e_.

### Molecular Exclusion from the Intercalated Disc

In order to estimate whether the different sized osmotic agents can permeate the intercalated disc (ID), the ID distribution of different sized fluorescently tagged dyes (0.5, 3, and 10 kDa) was measured by laser scanning confocal microscopy. Representative images in Figure 7A reveal the presence of the 0.5kDa dye in the ID, and the exclusion of the 3 and 10 kDa dyes. Figure 7B demonstrates how ID relative to bulk extracellular dye distribution was quantified by taking fluorescent intensity profiles of the dye through the Cx43 immunolabelled ID. By quantifying the intensity of dye beyond the Cx43 signal (lateral membrane) relative to the dye at the peak of the Cx43 signal, summary data on relative ID localization reveal that 0.5kDa dyes can permeate the ID with a normalized fluorescence intensity of 3.6 (95%CI: ±0.88), which is significantly greater than 1, indicating the dye accumulates in the ID. Relative normalized fluorescence intensity for 3kDa dye was 0.30 (95%CI: ±0.20) and 10kDa dye was 0.12 (95%CI: ±0.04), which are both significantly below 1, indicating these compounds were largely excluded from the ID. The data suggest that even if 66.5kDa albumin freely diffused into the bulk interstitium, it would not penetrate the ID, nor would dextran 70kDa or 2MDa. In short, small molecular weight molecules <3kDa can penetrate the ID and cause osmotically mediated dehiscence, while larger molecules may be size restricted from the ID and prevent or reduce fluid accumulation in these nanodomains.

**Figure 7.**
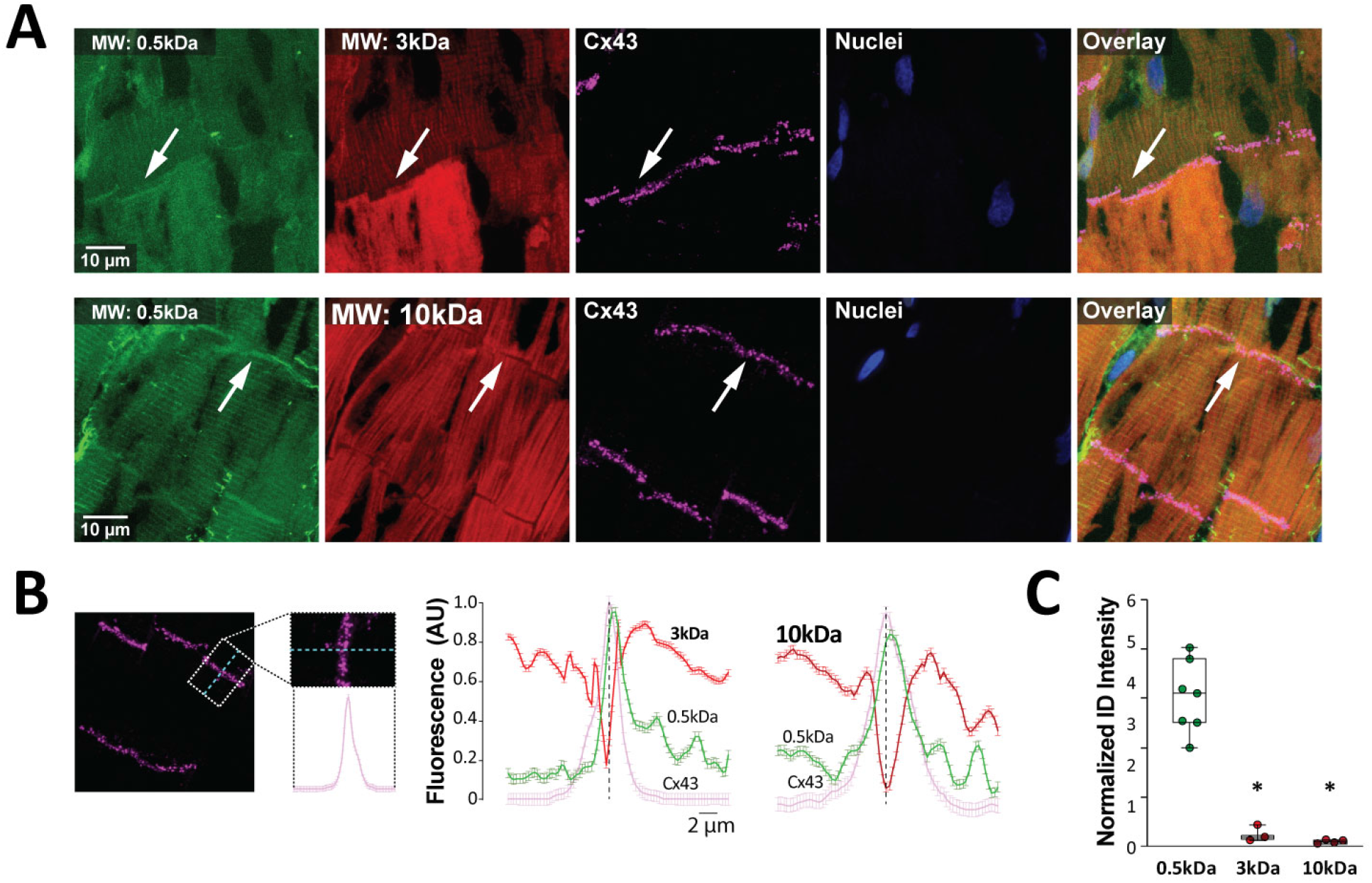
Molecules smaller than 0.5kDa can permeate the intercalated disc. **A.** Representative images of ventricular tissue slices perfused with 0.5, 3, and 10kDa fluorescent dyes. Panels demonstrate extracellular diffusion; Cx43 signal is used to identify the intercalated disc (ID), and nuclei are shown alone and with overlay of all signals. **B.** A region of interest is identified around the Cx43 ID signal and intensity of signal is averaged to identify the peak of the Cx43 signal. The same spatial averaging with 0.5, 3, and 10kDa signal demonstrates a dip in 3 and 10kDa signal with a peak in the 0.5kDa dye demonstrating the 0.5kDa dye permeates the ID, but the other two dyes do not. **C.** The intensity of the dye fluorescent signal at the peak of the Cx43 signal is normalized to the intensity of signal beyond the ID to reveal that 3 (n=3 images) and 10kDa (n=4 images) fluorescent signal are significantly lower than 0.5kDa (n=7 images) signal intensity. *p<0.05 relative to 0.5kDa, one way ANOVA with Dunnett’s correction.

### Ventricular Perinexus

To test the hypothesis that different sized osmotic agents differentially modulate ID volume, we quantified the width of the cardiac perinexus with transmission electron microscopy (TEM). Representative perinexi from the LV of hearts are shown in Figure 8A at 150,000X magnification. The osmotic agents differentially modulated cell membrane separation immediately adjacent to the GJ plaque within 15 minutes of osmotic agent perfusion. Specifically, and relative to Time Control, mannitol significantly increased W_P_ by 4.3nm (95%CI: ±3.1), albumin significantly narrowed W_P_ by -5.1nm (95%CI: ±2.9), dextran 70kDa did not significantly change W_P_ (Diff: 0.5nm, 95%CI: ±2.1), and dextran 2MDa significantly narrowed W_P_ by -3.9nm (95%CI: ±1.8) as shown in Figure 8B. Importantly, perinexal quantification revealed an inverse correlation between W_P_ and conduction. Specifically, mannitol *widened* W_P_ and *decreased* CV consistent with previous reports,^28^ and albumin and dextran 2MDa *narrowed* W_P_ and *increased* CV.

**Figure 8.**
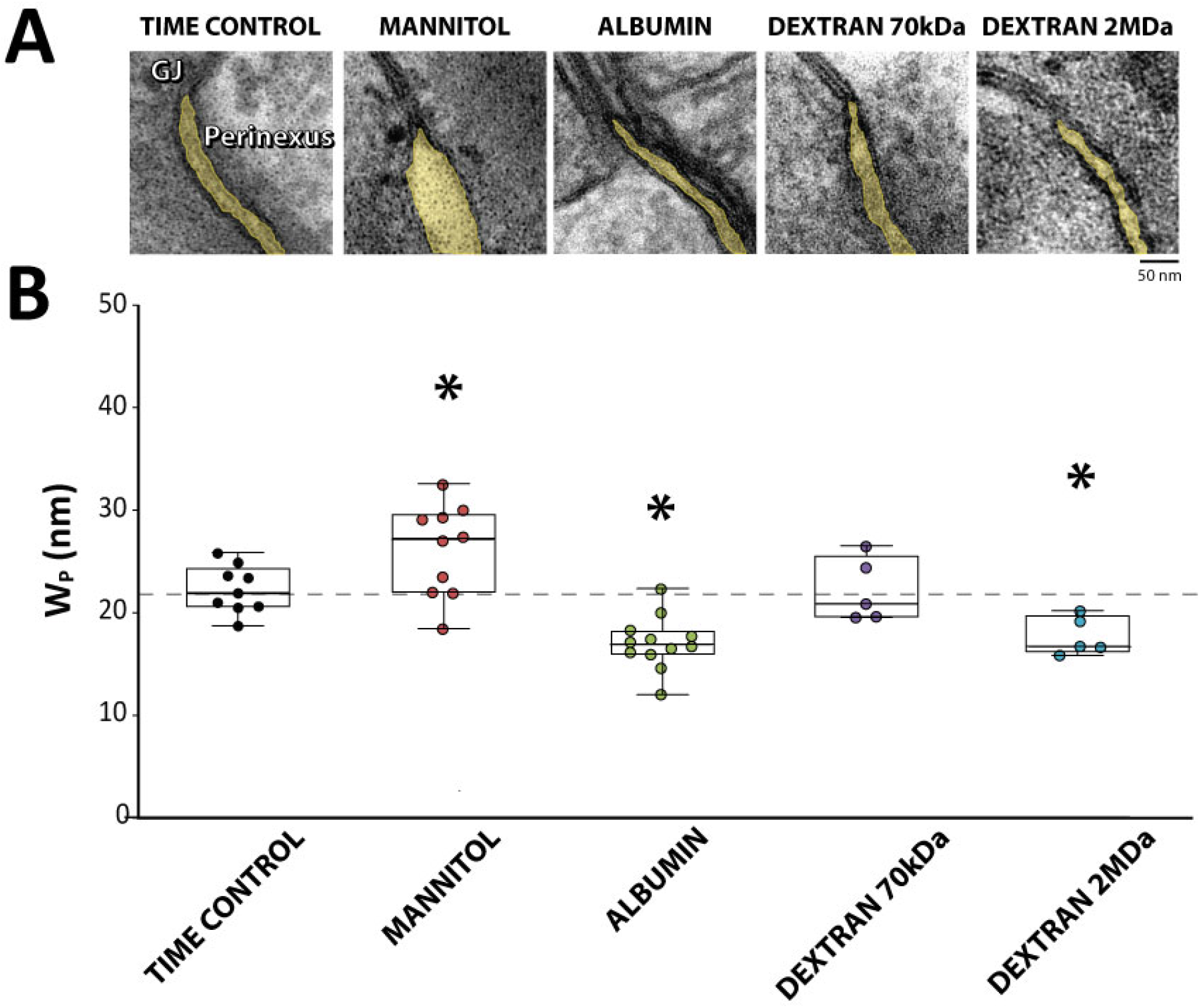
C*ardiac perinexal width inversely correlates with CV*. **A.** Representative TEM images of ventricular perinexi (shaded yellow) suggest osmotic agents differentially modulate perinexal width (W_P_). **B.** Summary data reveal that relative to Time Control (N=9 hearts, n=140 images), mannitol significantly increases (N=10 hearts, n=170 images), albumin significantly decreases (N=12, n=135 images), Dextran 70kDa does not change (N=5 hearts, n=74 images), and Dextran 2MDa significantly decreases (N=5 hearts, n=75 images) W_P_. Dashed line represents mean W_P_ for Time Control. *p<0.05, as evaluated with a generalized linear mixed effects model specifying a gamma distribution and identity link between samples collected from the same heart.

### Self-Attenuation

To this point, presented data demonstrate that CV is inversely correlated to W_P_. Yet, the relationship between W_P_ and CV is biphasic. For very narrow perinexi, CV can be directly proportional to W_P_ by altering sodium channel self-attenuation.^12, 35, 56^ In a final set of experiments, we hypothesized that combining dextran 70kDa with albumin would further decrease W_P_ and slow CV relative to albumin alone. Representative perinexal images and summary data in Figure 9A demonstrate that concurrent perfusion of albumin and dextran 70kDa significantly decreased W_P_ relative to albumin alone by 2.3nm (95%CI: ±2.0).

**Figure 9.**
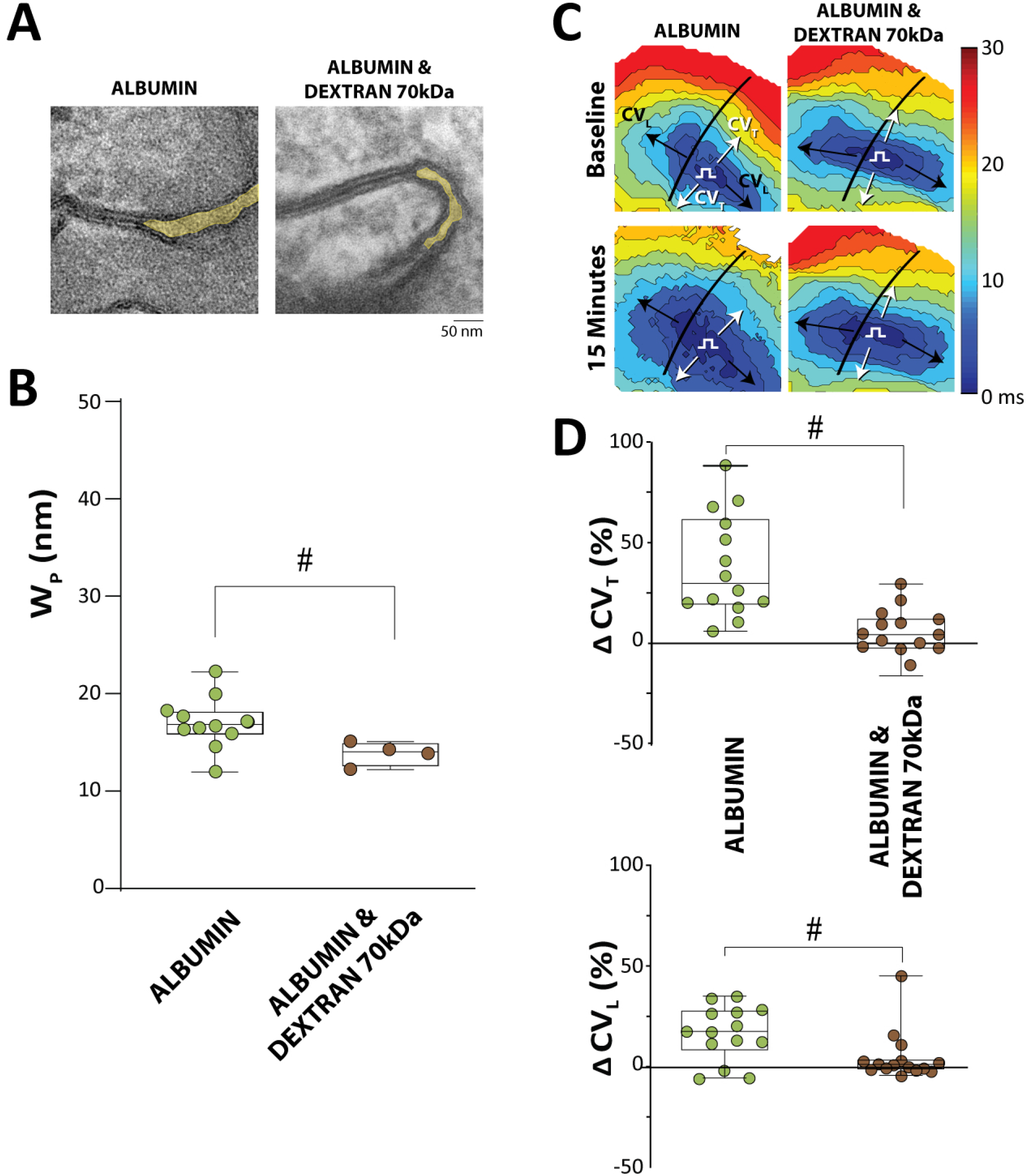
Dextran 70kDa in the presence of albumin can narrow perinexal width and slow conduction relative to albumin alone. **A.** Representative TEM images of ventricular perinexi (shaded yellow) and **B.** summary data reveal that simultaneous perfusion of albumin and Dextran 70kDa (N=4 hearts, n=60 images) significantly narrows perinexal width (W_P_) relative to albumin alone (N=10 hearts, n=150 images), #p<0.05 relative to albumin, as evaluated with a generalized linear mixed effects model specifying a gamma distribution and identity link between samples collected from the same heart. **C.** Representative activation time isochrones. Arrows indicate the conduction velocity directions of interest in longitudinal (CV_L_-Black) and transverse (CV_T_-White) propagation, and П notes the approximate site of pacing in the field of view. **D.** Summary data reveal that cardiac conduction is significantly slower in hearts treated with both albumin and Dextran 70kDa (N=15 hearts) relative to albumin alone (N=14 hearts). #p<0.05, unpaired two-tailed t-test. Note W_P_ data for albumin alone is replicated from figures 8 for comparison.

On average, albumin perfusion increased CV_T_ relative to baseline (Diff: 38%, 95%CI: ±13.3). Whereas the combination of albumin and dextran 70kDa did not significantly increase CV_T_ (Diff: 4.8%, 95%CI: ±11.8). Therefore, albumin and dextran 70kD slowed CV_T_ relative to albumin alone (Diff: 33.3%, 95%CI: ±14.9). On average, albumin perfusion increased CV_L_ relative to baseline (Diff: 16.2%, 95%CI: ±7.1). Whereas the combination of albumin and dextran 70kDa did not significantly increase CV_T_ (Diff: 4.5%, 95%CI: ±6.2). Therefore, albumin and dextran 70kD slowed CV_L_ relative to albumin alone, (Diff: 11.7%, 95%CI: ±9.8). The data demonstrate that despite increasing conduction on its own, dextran 70kDa slows conduction in the presence (Figure 9B) of albumin, potentially due to additive effects on perinexal narrowing leading to self-attenuation.

## DISCUSSION

Herein, we present two primary observations: 1) The cardiac perinexus, a specialized nanodomain of extracellular space within the intercalated disc (ID), can be osmotically expanded and collapsed on a timescale of less than 15 minutes, and 2) The regulation of cardiac epicardial conduction velocity (CV) is biphasically related with perinexal width (W_P_). Among the conditions tested, we did not observe significant changes in total Cx43 expression or phosphorylation at S368, nor did we find altered gap junction coupling (GJC) estimated by fluorescence recovery after photobleaching (FRAP). Additionally, the lack of a linear correlation between CV and altered tissue resistance argues for a minimal role in cable-like conduction mediated by gap junctions (GJ) alone. Furthermore, the osmotic agents did not significantly alter peak sodium current, although the change could have been below the resolution of detection. Our interpretation of these data is that extracellular volume changes in sodium channel-rich nanodomains such as the ID perinexus, rather than bulk extracellular resistance (Re), play a substantial role in modulating CV by an ephaptic mechanism in the absence of GJ remodeling.

In the present study, CV measurements for both mannitol and albumin were similar to those described earlier.^28^ Mannitol expanded W_P_ as we reported earlier.^19, 29^ Albumin and dextran 2MDa both decreased W_P_ and increased CV. When conduction changes were noted, they were primarily observed in the transverse direction of propagation. This is consistent with electrical wavefronts encountering more ID, with associated GJs and ephapses, along the short versus the long axis of cardiomyocytes for a given unit length. More specifically, as cardiomyocytes are highly polarized and IDs are located primarily on the edges of the long-axis, transverse propagation requires the electrical wavefront to “zig-zag” through the lateral edges. Since conduction across the ID is slower than through the myocyte,^57^ the increased number of junctions in the transverse, relative to the longitudinal direction will multiply the effect of modulating junctional communication on cardiac conduction. Interestingly, 40 g/l dextran 70kDa alone modestly but significantly increased CV in the whole-heart preparation, which seems to be in contrast to a previous study that found conduction slows with dextran 70kDa.^6^ Unlike our study, albumin was present in the baseline perfusion solution of the previous independent study. This being said, we replicated previous findings that dextran 70kDa in the presence of albumin will prevent the conduction increase associated with albumin alone. We propose the mechanism of CV slowing with dextran 70kDa and albumin is more dependent on perinexal narrowing resulting in ephaptic self-attenuation than a decrease in bulk extracellular impedance. Self-attenuation is predicted to occur when sodium channels decrease extracellular potential or sufficiently deplete extracellular sodium such that the inward driving force of this ion is reduced in ephaptic nanodomains. The results of the experiments with concurrent perfusion of albumin and dextran 70kDa are consistent with previous demonstrations of self-attenuation,^19, 35^ and reconciles conduction differences observed during osmotic stress by our group^28, 29, 31, 37^ and others.^6^

### Gap Junctions

Cx43-formed GJs and R_e_ are widely accepted as major determinants of cardiac conduction.^58–60^ As the half-life of cardiac Cx43 is on the order of 1-2 hours,^61^ and we observed CV changes at 15 minutes, it is not surprising that we do not measure a significant change in overall connexin protein expression. The further absence of associated changes in pCx43-S368 level is consistent with the lack of observable change in functional GJC estimated from FRAP assays. We previously demonstrated that Cx43 expression levels were not significantly changed by the same concentrations of albumin and mannitol perfused in this study.^28^ Though we cannot definitively rule out changes to GJ assembly or other Cx43 post-translational modifications, our data, combined with the short time scale of conduction changes, suggest that the primary mechanism for conduction changes may not be significantly attributed to GJC.

Most computational models of EpC and GJC do not demonstrate a biphasic relationship between CV and cleft width at “nominal” GJC,^12–15^ leading to the hypothesis that EpC is not particularly relevant during conditions of “normal” GJC. These models often reveal CV slowing secondary to widened cleft widths when GJC is reduced by more than 80%.^12–15^ As a result, a recurring question is whether EpC has a significant impact on cardiac conduction when GJC is nominal. In addressing this question, it is important to note that early computational models of cardiac conduction mediated by EpC and GJC often assumed normal GJ conductivity that is significantly greater^12–15, 57^ than has been measured experimentally between ventricular cardiomyocytes. Specifically, direct measurement of GJ conductances by dual-cell patch clamp yields estimates ranging from 3 to 2530 nS, with the majority of studies reporting an average GJC of less than 550nS, independent of species.^20–27^ Based on this consensus average, actual levels of GJC between myocytes are likely an order of magnitude smaller than the 2000nS values typically used as a nominal estimate of coupling in previous computational models. Furthermore, even in models that do not incorporate EpC, GJC is often tuned to lower values than those experimentally measured for the purpose of producing *in silico* CV values consistent with measurements made in heart.^62, 63^ It is possible that unaccounted for EpC could be a reason for non-physiologic GJC tuning that is necessary to replicate experimentally measured CV in hearts. Importantly, when one retrospectively re-evaluates these key early models in the context of consensus levels of GJC experimentally reported in the literature, EpC emerges as a major determinant of CV under normal, as well as pathological conditions of GJC.

### Perinexal Width

In addition to the aforementioned biphasic predictions of the CV-cleft width relationship in computational models of cardiac conduction, computational models that include both EpC and GJC also predict that CV sensitivity to GJC is reduced when intercellular separation is below approximately 20 nm and significantly enhanced at much wider cleft widths. These predictions have since been validated in mice and guinea pigs.^29, 35, 64^ Additionally, expansion of the perinexus has been associated with chronic atrial fibrillation in humans.^49^

Based on the observations that CV is inversely correlated to W_P_ with the individual osmotic agents, and that combining albumin and dextran 70kDa narrowed W_P_ more than albumin alone and prevented the albumin mediated CV increase, the data can be interpreted to support an ephaptic mechanism of osmotically regulated cardiac conduction. Specifically, very narrow perinexi are associated with relatively slow conduction (albumin and dextran 70kDa). Expanding the perinexus first increases CV (albumin or dextran 2MDa), peaks (dextran 70kDa) then starts to slow CV (control), and eventually dramatically slows CV as perinexi are severely disrupted (mannitol). These observations are consistent with an expected biphasic CV-W_P_ relationship predicted by models which include both EpC and GJC. ^11–14, 65^

While we frequently report W_P_ near 20nm, we also have demonstrated that the absolute value is dependent on the blinded experimentalist who traces the boundary of the perinexus in transmission electron microscopy images.^49^ Additionally, estimating a 3 dimensional volume from 2 dimensional images limits the precision of estimating perinexal volume, and therefore correlation between CV and W_P_. Even so, the relationship between CV and W_P_ is biphasic and therefore non-linear. Regardless, modelling and experimental data should be compared cautiously when attempting to establish numerical relationships between something as simple as channel conductance, whole-cell electrophysiology, and tissue level responses. For example, many previous models often assume a cylindrical geometry of myocytes with a surface area that significantly underestimates the tortuosity of the ID. Incorporating ion tracking^13^ or simple structural detail into the model such as interdigitation or the known disposition of larger GJs to encircle the ID^15^ modulates the response and range of CV observed with altered cleft width. The estimated CV values in EpC models are also dependent on sodium channel density in the ID,^15^ and the location of sodium channel clusters and their relative apposition within the ID.^18^ While it is tempting to infer a nominal starting point for any parameter on a predicted curve and speculate if CV should increase or decrease with a specific change in W_P_, this approach is likely premature. New studies utilizing models with enhanced structural detail, biophysics of spatial interactions, and ion channel clustering will likely yield important new insights into the effect of EpC on electrophysiology.^56, 66^

### Extracellular Impedance

The electrochemical impedance experiments revealed significant R_e_ changes at a similar time scale when conduction changes, as reported previously.^28^ Further, the data were collected on a similar temporal scale to the measurements of previous studies that carefully quantified tissue resistance at each point when conduction was measured.^6^ The physics of cable-like conduction through GJs alone is not compatible with our results, as R_e_ did not directly correlate with conduction. Notably, whereas both the 70kDa and 2MDa dextrans increased R_e_ and should have slowed conduction, they did not: Specifically, 70kDa and 2MDa dextran significantly increased CV. Furthermore, mannitol decreased R_e_ while simultaneously slowing CV as reported previously,^28^ but according to models only including GJC, mannitol should have increased CV.

Importantly, dextran 70kDa increased R_e_ by 21% in guinea pig whole-heart, similar to what was reported by others in isolated papillary muscle.^6^ To address the possibility that one or more osmotic agents altered GJC, cell size, membrane capacitance or any additional intercellular electrical parameters, we modeled the cellular pathway using a constant phase element (CPE). Tissue is composed of various cell types with different shapes and membrane properties, resulting in a distributed cell membrane time constant – the response speed of the cell membrane to electrical stimulation. This distributivity is what we quantified in the parameter, η. Since this parameter did not significantly change with Time Control, or with the addition of any of our osmotic agents, we conclude that the cell membrane time constant distribution on the tissue level was not substantially affected by our interventions. Therefore, the changes to R_e_ described above are likely a result of changes to the extracellular compartment, i.e. interstitial space. The lack of change in η is also consistent with non-significant changes induced by the osmotic agents in Cx43 expression or FRAP coupling assays. Importantly, macroscopic impedance measurement and its relationship to the cable theory is at best complicated and requires additional study.

### Sodium Channels

Despite the non-significant change in peak sodium current with osmotic agents, it is possible that conduction changes reported in this and previous studies^28, 30^ could be attributed to a change in peak sodium current below our resolution of detection. It should be noted here that the statistical approach may play a role in the ability to report a significant change in peak conductance. Specifically, if each current obtained from a cell is inappropriately considered an independent measurement, regardless of which heart it was obtained from, a simple non-paired Student’s t-test would suggest that mannitol modestly decreases peak sodium current on the order of 20%. If mannitol does significantly decrease peak sodium current, this and previous studies may over associate conduction slowing with perinexal widening. It is possible that a relatively small sodium channel inhibitory effect of mannitol exacerbates conduction slowing under these conditions as well. Yet, perinexal expansion associated with a peptide developed to disrupt sodium channel beta subunit trans-adhesion did not affect peak sodium current density and also slowed conduction in the same isolated guinea pig whole-heart preparation.^31^

The computationally predicted relationship of CV to W_P_ is also non-symmetric in the self-attenuation (positive correlation) and self-activation (negative correlation) phases of conduction. It is therefore difficult to know where the “normal” heart may reside on the CV-W_P_ curve because this relationship is also modulated by factors such as sodium and potassium channel localization to the ID, channel unitary conductances, channel availability, cell size, and extracellular sodium and potassium.^12–15, 19, 67–70^ Importantly, we provide experimental evidence that the degree of perinexal narrowing or expansion is associated with different magnitudes of CV increase or decrease, respectively. Thus, at present, it is difficult to state to what degree the different osmotic agents should change CV for a given change in W_P_. We can say with confidence that the change in CV with narrower and wider W_P_ is not uniform. Regardless, evidence continues to support the conclusion that cardiac conduction is well-described by models including both EpC and GJC.

### Membrane and Intracellular Pathways

The 4-electrode distributivity measurement, a metric of cell membrane capacitance, cytosolic resistance and GJ resistance, does not significantly differ between controls and cells or tissue incubated with the osmotic agents. This inconclusive result does not support nor exclude the possibility that osmotically induced changes in cellular parameters are significant determinants of CV. The lack of distributivity change is inconsistent with previous results when cell volume was measured by cell morphometry for example. Specifically, we and others reported that mannitol can decrease *isolated* ventricular myocyte volume by approximately 7% to 25% for concentrations of mannitol between 50 and 143mM.^28, 71, 72^ The discrepancy and lack of significant differences in metrics of intracellular impedance in this study could therefore be a result of: a change that is below the detection resolution of the technique, intra and interventricular heterogeneity,^73^ inter-animal variability, and/or variability secondary to cellular isolation.^74^ Importantly, a previous computational model of conduction found that the effects of cell size and CV are complexly dependent on both forms of electrical coupling.^16, 70^ Specifically, with relatively high GJC (2500nS), decreasing cell size can slow conduction, and EpC can modestly steepen that relationship. In contrast, with relatively nominal but closer to normal GJC (100nS), decreasing cell size can increase CV, but once again EpC modestly increases the steepness of this relationship.^70^ Since GJC has been measured over a large range of values, and the average value in literature is closer to 550nS,^20–27^ we do not know which result to expect. Therefore, the effect of parameters such as cell volume on the observed changes in CV still requires significantly more independent investigation for a number of reasons including: 1) Morphometric changes in myocyte geometry are likely different in an isolated, unloaded cell relative to cells in intact myocardium. 2) The osmotic agents will partition differently between vascular, extracellular, and ID compartments based on osmolyte permeability and size. 3) The molarity of the osmotic agents used in this study are not identical.

### Limitations

No intervention can be applied without experimental consequence. Osmotic agents employed in this study have well-established effects in addition to changing oncotic pressure, and some are used clinically for these and other reasons.

*Albumin*, for example, has been used to regulate blood pressure due to its binding affinity to the vascular endothelium, leaving vessels at a rate of approximately 5% per hour. Albumin furthermore regulates vascular function and can bind to a number of cations, fatty acids and hormones, in addition to acting as an antioxidant.^75^

*Mannitol*, a sugar alcohol, has been used as a hyperosmotic agent to treat high intracranial pressure.^76, 77^ With appropriate dosing and time courses of treatment, mannitol will decrease interstitial edema by drawing fluid out of the cerebral parenchyma; though at higher doses and over long time courses, mannitol will also cross the blood-brain barrier. Once across the blood-brain barrier, mannitol will then have the opposite intended effect, more similar to what we observe in cardiac tissue: drawing fluid into tissue and exacerbating edema.

*Dextrans* of various sizes have been used, like albumin, as a blood volume expander, with larger dextrans having a more pronounced effect on blood viscosity. Dextran 70kDa has antithrombotic properties and reduces the inflammatory response and troponin-I release after cardiac surgery.^78, 79^ It remains unknown, however, whether the other non-osmotic properties affected our results over the much more acute time frame of less than 15 minutes studied here.

Despite our efforts to select osmotic agents with minimal binding or metabolic effects, we cannot definitively exclude any such effects and advise the reader to cautiously interpret our results about any implications for clinical treatment.

As mentioned previously, experiments were conducted in excised, isolated, male retired breeder, guinea pig ventricles that were perfused with di-4-ANEPPS and preserved for hematoxylin and eosin (H&E) analysis and electron microscopy with formalin and glutaraldehyde, respectively. As such, all values of structural dimensions presented – e.g. W_P_ and interstitial volume (VIS) – should be interpreted without implication of explicit *in vivo* values. Additionally, while our CV results are similar to those published previously,^28^ we observed no significant differences in interstitial volumes between interventions, though all of our values were similar to the controls reported previously. It is worth noting that the reported interstitial volume differences may be attributed to a shorter protocol (15 minutes in this study vs 60 minutes in previous studies), different fixatives (formalin vs glutaraldehyde), different analysis programs (positive-pixel vs color deconvolution), and the statistical approach. Interestingly, previous work suggests that 30 minutes were required for changes in R_e_ to reach steady-state.^6^

Furthermore, albumin is difficult to work with in optical mapping experiments, because signal quality degrades quickly as demonstrated previously^30^ and in Figure 2A. The reasons for this are not entirely understood since the absorption peak of albumin is approximately 280 nm^80, 81^ and peak excitation for di-4-ANNEPS is near 480 nm.

Optical mapping also provides fluorescent signals from a volume of tissue that includes intramural cells, and volume averaging may “blur” the upstrokes of optical action potentials and captures transmural propagation patterns.^82–86^ The values of conduction between the left and right ventricle may also be affected by transmural rotational anisotropy given the difference in myocardial thickness.^87^ Therefore, caution should be exercised when considering the absolute CV values presented herein. However, our group is careful about conducting positive and Time Control measurements with optical mapping under a variety of conditions previously reported to affect conduction: sodium channel inhibitors,^29, 88^ GJ inhibition and Cx43 genetic loss of function^35, 69^ mannitol or albumin,^28, 30^ and changes in extracellular potassium.^69^

With regards to whether edema confounds epicardial measures of CV due to volume averaging, we currently discount this hypothesis for a few reasons. First, dehydrating tissues should bring more cells into the field of view epicardially and transmurally. Epicardial recruitment of cells into an imaging volume should decrease the speed of conduction by increasing the number of IDs in each pixel, even if the dimmer mid myocardium would increase conduction. The epicardial recruitment effect is inconsistent with our finding that solutions associated with increased or decreased R_e_, a metric of bulk hydration, do not correlate with the directional change in conduction. Furthermore, we provide confirmatory evidence that dextran 70kDa in the presence of albumin is associated with slower conduction when compared to hearts perfused only with albumin, and this is consistent with studies using microelectrodes, the same osmotic agents, and papillary muscles that lack rotational anisotropy.^6^

Finally, within our tissue impedance experiments, electrode array orientation was visually aligned with local fiber orientation. However, as fiber orientation changes in 3 dimensional myocardial tissue, we cannot compare tissue impedance changes between experiments, only between interventions in the same heart. As a result, only the finding that R_e_ changed has practical meaning, but the absolute values of R_e_ and magnitude of change do not have direct physiologic relevance. Additionally, the measurement of R_e_ does not distinguish between bulk extracellular volume expansion or changes to conductivity in the ID. Both volume changes could change R_e_ in the same direction.^89, 90^ However, our observations that CV does not change proportionally in response to R_e_ argues against bulk extracellular volume expansion as the principal determinant of CV in these experiments.

### Conclusions

Clinically relevant osmotic agents may impact cardiac conduction by a previously unappreciated mechanism of modulating nanodomain separation between cardiomyocytes in the perinexi of ID. Specifically, this study demonstrates that expanding ID nanodomains densely expressing ion handling proteins can both increase and decrease cardiac conduction consistent with computational predictions of cardiac conduction dependent on both EpC and GJC. While we cannot definitively exclude the possibility of off-target effects from the experimental interventions studied, our data suggest that EpC modulation of one type of ID nanodomain, the perinexus, is a major determinant of cardiac conduction. The perinexus therefore continues to emerge as a therapeutic target for managing cardiac arrhythmias, and further research is needed to better understand the conditions when perinexal expansion represents a beneficial adaptation or pathologic remodeling.

